# Sequence learning in a single trial: a spiking neurons model based on hippocampal circuitry

**DOI:** 10.1101/2020.01.09.898064

**Authors:** S. Coppolino, M. Migliore

## Abstract

In contrast with our everyday experience using brain circuits, it can take a prohibitively long time to train a computational system to produce the correct sequence of outputs in the presence of a series of inputs. This suggests that something important is missing in the way in which models are trying to reproduce basic cognitive functions. In this work, we introduce a new neuronal network architecture that is able to learn, in a single trial, an arbitrary long sequence of any known objects. The key point of the model is the explicit use of mechanisms and circuitry observed in the hippocampus, which allow the model to reach a level of efficiency and accuracy that, to the best of our knowledge, is not possible with abstract network implementations. By directly following the natural system’s layout and circuitry, this type of implementation has the additional advantage that the results can be more easily compared to experimental data, allowing a deeper and more direct understanding of the mechanisms underlying cognitive functions and dysfunctions, and opening the way to a new generation of learning architectures.

## Introduction

If we decide to have a second look at a nice and unique piece of art, while navigating through the halls of a museum that we are visiting for the first time, we do not need a long and extensive learning session. Just a single walk during the initial visit will usually (although not always) be sufficient to reach our goal. This is in striking contrast with the notoriously long and difficult process that any deep learning algorithm (reviewed in Schmidhuber, 2015) needs to follow to learn a trajectory through a sequence of objects. Depending on the problem, training a neural network to produce the correct sequence of outputs in the presence of a series of inputs can take a prohibitively long training time. This is a clear indication that something important is missing in the way in which computational models try to reproduce basic cognitive functions. Nevertheless, using customized network architectures and clever learning algorithms, recurrent neural networks such as Long Short-Term Memory or Bidirectional Recurrent neural networks have reached impressive practical results (reviewed in Yu et al., 2019). These networks, however, have little resemblance with the biological architecture of the hippocampus, a brain region well known to integrate space, time and memory (reviewed in Eichenbaum, 2017), and to also deal in an extremely efficient way with the task to learn how to follow cues during spatial navigation (in a museum, a maze, a series of roads, a building, etc.). In this work, we tried a new approach to learning a sequence of objects, by building a neuronal network architecture that significantly departs from the current mainstream of deep learning algorithms; we show that by assembling cell and circuit properties observed in the hippocampus, the resulting network is able to learn an arbitrarily long sequence of arbitrary objects in a single trial, opening the way to a new generation of artificial intelligence algorithms.

## Methods

The network was implemented in PYNN (Davison et al., 2009), and it was composed from 21 leaky integrate-and-fire (LIF) neurons with the same electrophysiological properties. During the learning phase, the synaptic weights evolved following a spike-time-dependent synaptic plasticity rule (STDP) implemented by considering experimental findings in the hippocampus (Nishiyama et al., 2000). The Robot Operating System (ROS, https://www.ros.org/about-ros/) was used to build a basic closed-loop environment to get input and direct the movement of a virtual robot. Several python custom transfer functions were also implemented to integrate the PYNN and ROS code into the NeuroRobotics Platform of the Human Brain Project (NRP-HBP, https://www.humanbrainproject.eu/en/robots/). As soon as the corresponding paper is published, all model and simulation files used for this work will be publicly available on ModelDB (https://senselab.med.yale.edu/ModelDB/), and on the HBP Brain Simulation (https://www.humanbrainproject.eu/en/brain-simulation/brain-simulation-platform/) and NeuroRobotics (https://www.humanbrainproject.eu/en/robots/) Platforms.

## Results

For the purposes of this work, the network was composed of 21 LIF neurons arranged as a feed-forward network schematically shown in Fig.1. According to their location and connectivity in the network, the different neurons were labeled as Place Cells (*PC*), object cells (*obj*), persistent firing cells (*PF*), and left and right cells (*L* and *R*, respectively). It should be stressed that we used cell and circuit properties observed in the hippocampus, such as:

1. cells selectively firing in presence of specific inputs associated with a given object; this sparse and explicit coding has been observed experimentally (e.g. Quiroga et al., 2005), and we have previously shown how it can be implemented in a biophysically detailed model (Migliore et al., 2008). We indicated these cells as *obj* in Fig.1. In general, by *object* we mean a set of incoming fibers collectively coding for a number of features forming a sparse representation of a multisensorial input, and targeting a single or very few neurons. Each *obj* cell makes synaptic contacts with all *PC* cells. For the sake of simplicity, in this work we assumed that only one *obj* cell will be activated when the corresponding object is presented as input.
2. Persistent firing cells, *PF* in Fig.1. These cells are active during the maintenance period of cognitive tasks (Boran et al., 2019), and imply internal processing in the absence of an external input. We assumed that they are activated by an input object, and make synaptic contacts with all *obj* cells except the one corresponding to the same object, as illustrated by the grey circles between *PF* and *obj* cells in Fig.1. Consistently with experimental findings (Boran et al., 2019), a *PF* cell begins tonic firing just before the corresponding *obj* cell becomes inactive, this firing ends when new object is presented as input.
3. Place Cells, *PC* in Fig.1. A hallmark of spatial navigation, these cells are commonly found in the CA1 hippocampal region (reviewed in Danjo, 2019); they fire when an animal is in a specific spatial location, and do not represent just space but also contextual information (Gulli et al., 2020). In our network, they are responsible for generating a signal when an object is observed with a specific head direction (HD); this output is used to control robot movement.
4. Avoidance path. This is another mechanism commonly observed in hippocampal CA1 area, that also depends on the integrated activity with other brain areas (reviewed in Giovannini et al., 2015); in the network, we implemented this signal by generating activity in the left/right direction, opposite to the location of unknown objects such as walls and obstacles.

**Fig. 1:**
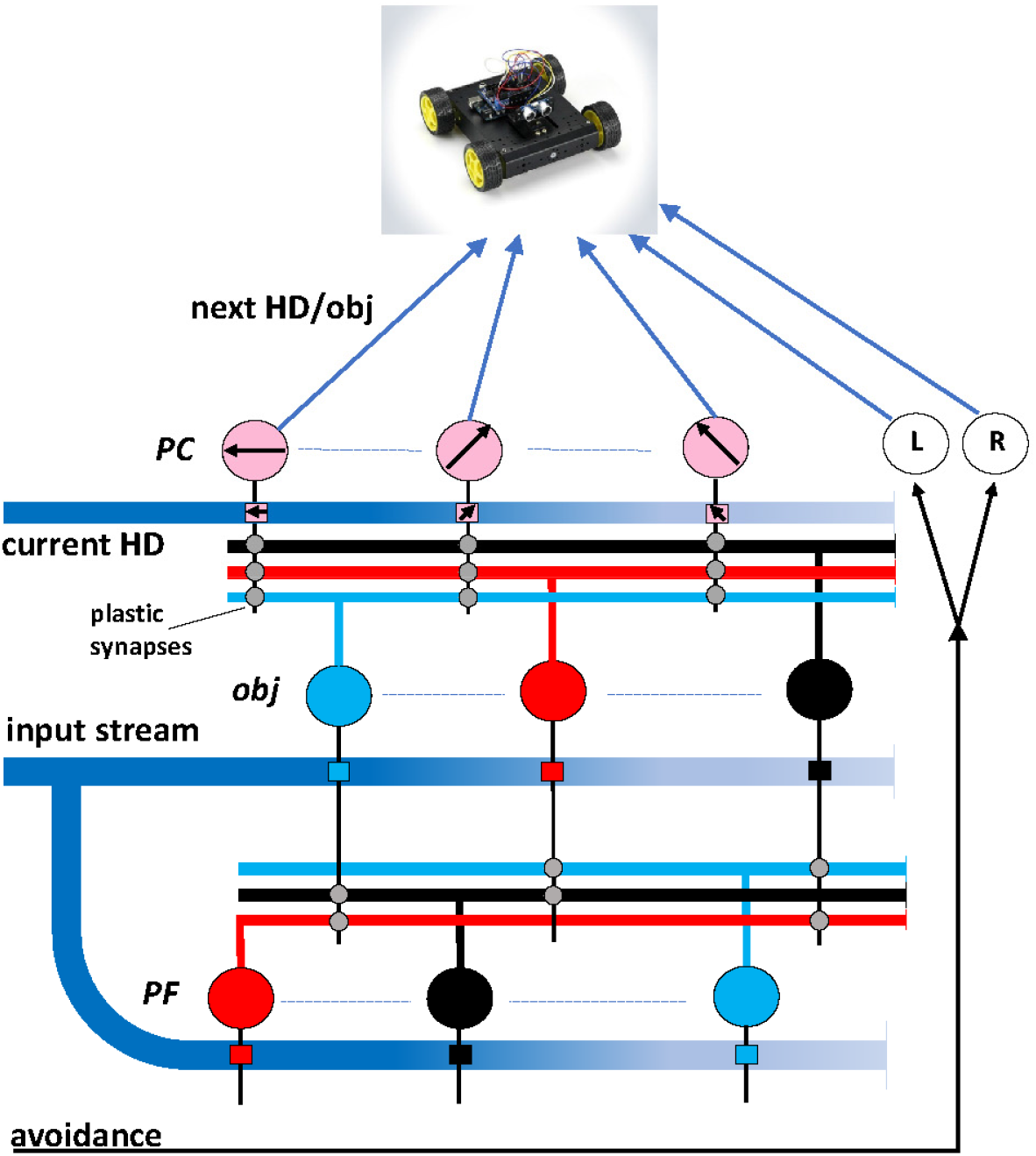
Network architecture. Schematic representation of the network. Circles of different colors (blue, red, and black) represent neurons tuned to different input cues (different objects in this case); Blue pathways represent external inputs carrying contextual information (object identification and the current Head Direction (HD), in this case); colored squares indicate that the relative cell will be activated whenever the corresponding object is present in the input stream. Grey circles indicate plastic synapses; obj, object cell; PC, Place cell; PF, persistent firing cell; L, left; R, right; For clarity, only 3 of the 12 PC cells used in the network are shown in the diagram.

There are two main inputs to the network: one representing the current head direction and one representing an object (large blue pathways in Fig.1). According to their content, these input pathways will activate one *obj*, one *PC*, and one *PF* cell, as represented by the small colored squares along both pathways. This is a simplified organization, to illustrate the proof of principle. The same general network organization will also work in a more natural layout, where a complex input will activate a number of *obj* neurons in a more distributed fashion.

We tested the network by connecting it to a basic virtual mobile robot, available in the NRP-HBP. A movie of the (single) learning trial and of recall tests, under different conditions, is presented as Supplementary material. The network dynamics and synaptic weights evolution during the exploration of an environment with three objects is illustrated in Fig.2. We assumed that these objects (screens of different colors in this case) are already known to the network, and therefore, that their presence in the input will generate a train of spikes in the corresponding *obj* cell.

**Fig. 2:**
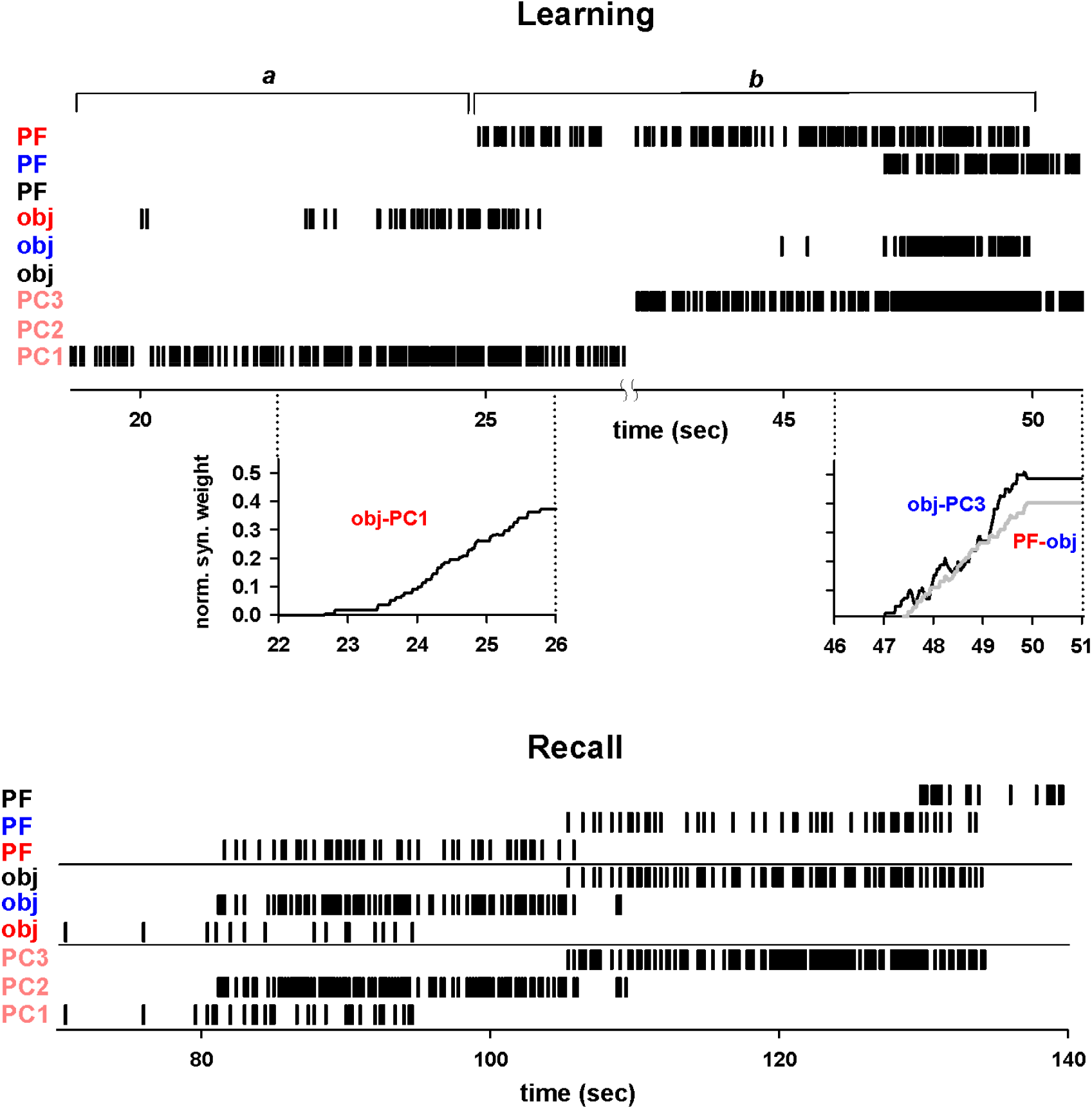
Firing patterns and synaptic weights evolution during learning and recall. (Learning) firing patterns during a typical learning session; phase a) and b) are described in the main text. PF and obj labels are color coded to represent the corresponding object. PC1-3 represent neurons originally tuned to fire for three different head directions; during the learning phase, two of them (PC1 and PC3) will become place cells, and will signal to the robot the direction to take to go to the next object in the sequence. The insets at 22-26 and 45-51 sec show the evolution of selected synaptic weights. (Recall) raster plots of the same neurons in Learning during a recall session.

The raster plots in Fig.2 represent the firing patterns of different cells during the learning and recall phases; they will be better appreciated if compared with the movie (see Supplementary movie).

### Learning phase

a. In the absence of a known input, the robot does a stereotypical space exploration by looking around counterclockwise to find a known object. This movement generates the sequential activation of the PC cells tuned to the different head directions (for clarity, this part is not reported in Fig.2). As soon as a known object enters the visual field (a red object in this example) it will activate the red *obj* cell, and its spikes will propagate to all *PC* cells; however, at any given instant, only one of the *PC* cells is also independently firing in response to the current head direction information (*PC1* in this example); the two concurring inputs on this neuron will potentiate the *red obj*-*PC1* synaptic weight (Fig.2, *learning*, inset at sec 22-26).
b. In the meanwhile, the *PFred* cell will also begin to generate spikes, which will be propagated to all the other *obj* cells, while the robot looks for the next object; when it sees it (a blue object, see Fig.2 *learning*, beginning at sec 45), two events occur: 1) the potentiation of the *PFred-blue obj* synaptic contact (Fig.2, inset at 46 sec, grey line), because of the concurrent activation of the *blue* and *PFred* neurons and, 2) the potentiation of the *blue obj–PC3* synapse (Fig.2 inset at 46-51 sec, black line), because of the concurrent activation of the *blue* and PC3 neuron (corresponding to the current head direction); this is a crucial point, and it is the basic process through which the network learns the direction to be taken after any given object in a sequence.
c. Points a) and b) are repeated until the goal is reached.

These steps, executed in a single pass until the goal is reached, conclude the learning phase. From the previous discussion, it should be clear that the network will correctly learn sequences of any length and any object combination; the key point is that the STDP mechanism is sufficient to potentiate the correct synapses to learn the direction to the next object.

### Recall phase

During a recall phase (see Fig.2, *recall*), the robot, controlled by the firing generated by the *PC* cells with potentiated synapses, will now follow a smooth path through all the objects (see Supplementary movie). Note that all neurons are sequentially activated following the learned sequence. This corresponds to the typical firing behavior observed experimentally for place cells during spatial navigation (e.g. Haimerl et al., 2019). In a different recall session (see Supplementary movie), we demonstrate how the robot is also able to avoid walls and obstacles along the path, through the activity of the avoidance cells (not shown in Fig.2).

## Discussion

In this work, we have introduced a new neuronal network architecture that is able to learn, in a single trial, an arbitrary long sequence of any known objects. The key point to obtain this result was the explicit use of mechanisms and circuitry observed in the hippocampus, instead of the usual artificial network of identical point neurons recurrently connected all-to-all and evolving following abstract learning rules. More generally, the model supports the idea that, by explicitly including biologically plausible interactions among the different neuron types in different cortical and subcortical structures (reviewed in Hinman et al., 2018), one can obtain a network able to perform as efficiently as in the real system. The crucial point is the ability to rapidly configure the network, following the type of conjunctive input processing experimentally observed at cellular level (Bittner et al., 2015). At this moment, this is the only model of spatial navigation able to produce this type of result. The main limitations of this simple proof of principle implementation are that:

a. the network does not learn new objects; this mechanism can be added to the network through a heterosynaptic competition process (Fiete et al., 2010; Rajan et al., 2016) that, in the context of our model, will be able to learn a new object and eventually include it in the network on the fly, during the learning phase.
b. confusion may arise in case of a sequence with multiple identical choices, such as when the same object appears in the sequence in two or more different places after a given object, and when it is also seen from the same angle; however, even a human could miss the right choice in this case, without additional cues or previous learning phases.
c. complex scenes, such as those including multiple known objects and contextual cues, cannot be properly processed with the current implementation; they require a mechanism binding together objects and context information, known to be in effect in the hippocampus to represent a scene (see e.g. Gulli et al., 2020); this aspect was outside the scope of this work.

Most, if not all, of these limitations can be taken into account by an inhibitory network, exerting its disambiguation action through lateral and feedback inhibition mechanisms (see e.g. Priebe and Ferster, 2008; Yu et al., 2014). A simple comparison with a typical recurrent neural network architecture, should make it clear that there is a striking difference with other model networks. Consider the simplest case of learning just one sequence, composed from *k* objects randomly chosen from a set of *n* different objects, without repetitions. The goal is to obtain a network able to generate the correct list of objects, starting from any point in the sequence. During a training session with an artificial network, the sequence must be presented many times in the input, to let the weights evolve until the error is small enough to ensure a good performance during a recall. The number of times will strongly depend on the length of the sequence and the details of the learning rule, and the accuracy cannot be guaranteed to converge to a good value. Instead, our network will learn the sequence in a single presentation, with 100% accuracy, no matter how long the sequence is.

In this work, we have demonstrated how the explicit and direct use of biological circuitry to model a cognitive function can reach a level of efficiency and accuracy that is not possible with abstract network implementations. In spite of its current limitations, it should be clear why this approach can lead to a new generation of artificial intelligence algorithms for spatial navigation. The network self-organizes to the correct functional state without any need for complex learning rules or conditional multinomial probability distribution functions keeping track of the different states visited during navigation. By directly following the natural system’s layout and circuitry, this type of implementation has the additional advantage that the results can be more easily compared to experimental data, allowing a deeper and more direct understanding of the mechanisms underlying cognitive functions and dysfunctions.

## Acknowledgements

This work was supported by the European Union’s Horizon 2020 Framework Programme for Research and Innovation under the Specific Grant Agreement 785907 (Human Brain Project SGA2). Editorial support was provided by Annemieke Michels of the Human Brain Project.

## Author contributions

SC implemented the model and performed all analyses; MM conceived the study and wrote the manuscript.

